# Genomic prediction of growth in a commercially, recreationally, and culturally important marine resource, the Australian snapper (*Chrysophrys auratus*)

**DOI:** 10.1101/2021.09.02.458800

**Authors:** Jonathan Sandoval-Castillo, Luciano B. Beheregaray, Maren Wellenreuther

## Abstract

Growth is one of the most important traits of an organism. For exploited species, this trait has ecological and evolutionary consequences as well as economical and conservation significance. Rapid changes in growth rate associated with anthropogenic stressors have been reported for several marine fishes, but little is known about the genetic basis of growth traits in teleosts. We used reduced genome representation data and genome-wide association approaches to identify growth-related genetic variation in the commercially, recreationally, and culturally important Australian snapper (*Chrysophrys auratus*, Sparidae). Based on 17,490 high-quality SNPs and 363 individuals representing extreme growth phenotypes from 15,000 fish of the same age and reared under identical conditions in a sea pen, we identified 100 unique candidates that were annotated to 51 proteins. We documented a complex polygenic nature of growth in the species that included several loci with small effects and a few loci with larger effects. Overall heritability was high (75.7%), reflected in the high accuracy of the genomic prediction for the phenotype (small vs large). Although the SNPs were distributed across the genome, most candidates (60%) clustered on chromosome 16, which also explains the largest proportion of heritability (16.4%). This study demonstrates that reduced genome representation SNPs and the right bioinformatic tools provide a cost-efficient approach to identify growth-related loci and to describe genomic architectures of complex quantitative traits. Our results help to inform captive aquaculture breeding programmes and are of relevance to monitor growth-related evolutionary shifts in wild populations in response to anthropogenic pressures.

## Introduction

Body size is considered one of the most important organismal traits because it influences several biological characteristics, from survivorship to fecundity (Peters 1986). It has significant ecological and evolutionary consequences, not just for the individual but for its population and community (Lorenzen 2016; Dijoux and Boukal 2021). Moreover, the somatic growth patterns of exploited fishes have a considerable role in the sustainability and economic viability of their fisheries stocks and aquaculture programmes (Gjedrem *et al*. 2012; Murua *et al*. 2017; Barneche *et al*. 2018). These characteristics make growth and related traits major targets for selective breeding and human-induced evolution studies (Sharpe and Hendry 2009; Gjedrem *et al*. 2012; Denechaud *et al*. 2020; Huang *et al*. 2021). Teleost fish typically show indeterminate growth, but both the growth rate and growth-related plasticity are determined by the genetic makeup and the prevalent environmental conditions (De-Santis and Jerry 2007; Boltaña *et al*. 2017; Tung and Levin 2020). However, despite the general importance of growth, the genetic bases of growth traits in marine species are still not well understood.

Anthropogenic activities are rapidly transforming environments, resulting in changes to the selection landscape that organisms to which are exposed, and consequently, phenotypic changes in response to altered selection pressures (Hendry *et al*. 2017; Laufkötter *et al*. 2020; Silvy *et al*. 2020). Fish growth rates can respond rapidly to external selective pressures by phenotypic plasticity or by changes in genotypic frequencies in the population (Bowles *et al*. 2020; Denechaud *et al*. 2020; Oke *et al*. 2020; Pinsky *et al*. 2021). Fish growth is also a quantitative trait that is easy to measure, and as such, can be used to assess physiological and demographic responses to human-mediated environmental changes (Denechaud *et al*. 2020). Climate change has increasingly and profoundly threatened fish biodiversity (Free *et al*. 2020; Huang *et al*. 2021). In fishes, changes in growth are the most direct and frequent responses to climate change (Rountrey *et al*. 2014; Heather *et al*. 2018; Ikpewe *et al*. 2021). In addition, pressures from capture fisheries have produced one of the fastest rates of phenotypic change observed in wild populations (Oke *et al*. 2020), with effects not only to targeted species but also to associated communities (Andersen *et al*. 2016; Dijoux and Boukal 2021). However, size-selective harvesting and climatic effects could be species and system specific (Denechaud *et al*. 2020), and despite increasing concerns, the evolutionary consequences have so far been investigated in only a limited number of species. Thus, the extent to which changes to growth rate in response to human activities are evolutionary or plastic, and the degree to which are they reversible, remain debated (e.g. Pinsky *et al*. 2021).

The idea of rapid evolutionary responses due to fisheries pressures and aquaculture practices is supported by both theoretical and empirical evidence. In theory, high rates of harvesting of a population will favour earlier sexual maturity and slower growth (Uusi-Heikkilä *et al*. 2015; Monk *et al*. 2021). It is also expected that selective breeding programmes increase the frequency of genetic variants underlying improved growth through the successive selective breeding of only the fastest growing individuals in each cohort. Experiments in the laboratory, where either large or small individuals are removed for several generations, have shown drastic changes in allele frequencies, loss of genetic diversity, and increased linkage disequilibrium in specific parts of the genome (Le Rouzic *et al*. 2020; Valenza-Troubat *et al*. 2021). Some genes and genomic regions associated with growth have been identified in a few marine fish species (Wang *et al*. 2015; Robledo *et al*. 2016; Wu *et al*. 2019; Zhou *et al*. 2019; Yang *et al*. 2020; Valenza-Troubat *et al*. 2021). However, because of the apparently polygenic nature and high phenotypic plasticity associated with the trait, the genetic basis of growth is not well understood, and its study is a persistent challenge (Wellenreuther and Hansson 2016; Gong *et al*. 2021; Valenza-Troubat *et al*. 2021). The combination of genotyping based on reduced genome representation datasets and genome-wide association studies (GWAS) potentially provides a cost-effective and convenient approach for accurate localization and identification of growth-related genomic regions in fishes.

The Australian snapper (*Chrysophrys auratus*) is a large (~1 m) and long-lived (>50 years) marine teleost (Fig. 1) that supports important recreational and commercial fisheries in Australia, New Zealand, and some Pacific Islands (Parsons *et al*. 2014; Cartwright *et al*. 2021). The species is also of major cultural importance, having been caught by New Zealand’s indigenous Māori people for hundreds of years, and is considered taonga (a treasure) by them. The species stocks have large biomass, but since the early 2000s most have shown declines, with some losing over 75% of their biomass (Cartwright *et al*. 2021). These reductions in snapper productivity have negative economic and ecological consequences that could be amplified by global climatic change (Ogier *et al*. 2020; Parsons *et al*. 2020). A strategy for reducing these negative effects, while enhancing wild snapper stock productivity, is the development of captive production. Although there is not yet commercial aquaculture of the species, research programmes intending to develop Australian snapper into a profitable and sustainable aquaculture species exist in New Zealand and Australia (Ashton *et al*. 2019a; Catanach *et al*. 2019). The goal of our work is to contribute to both the long-term New Zealand efforts to develop this aquaculture activity, and the understanding of the genetic bases of growth-related traits in marine fish more generally. We took advantage of several resources available from the New Zealand snapper aquaculture programme, including a chromosome-scale assembly (Catanach *et al*. 2019) and fish from the New Zealand snapper breeding programme representing extreme growth phenotypes following a grow-out phase in a seapen. Using these resources, we then applied reduced genome representation to conduct GWAS analyses to identify growth-related genetic variation in a cost-efficient way. In addition, we implemented a Bayesian mixture model to describe the genetic architecture of growth and to test the accuracy of size phenotype prediction based on single nucleotide polymorphism (SNP) genotypes. This information will help to improve the selection of high-quality broodstock and the monitoring of human-induced evolution in wild stocks.

**Figure 1.**
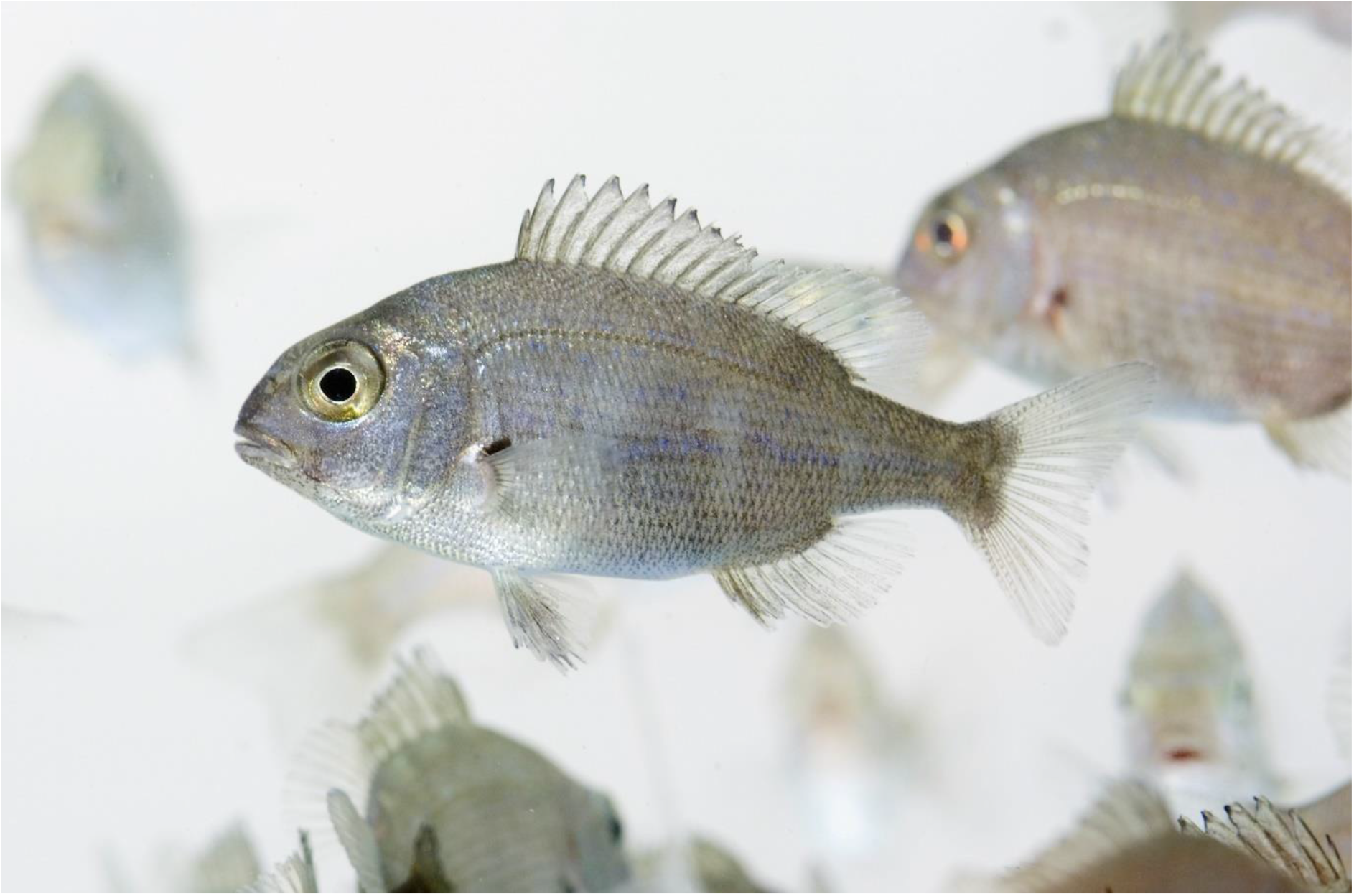
Juveniles of Australian snapper (*Chrysophrys auratus*) reared in captivity at The New Zealand Institute for Plant & Food Research Limited finfish facility.

## Methods

### Fish rearing and grow-out conditions in the sea pen

In 2016, a snapper breeding programme was started in Nelson (New Zealand) with wild-caught animals at The New Zealand Institute for Plant and Food Research Limited (PFR). Uncontrolled mass spawning of a wild-caught broodstock at the PFR finfish facility in Nelson produced two batches of eggs that hatched on 31 January and 7 February 2015, respectively. These F_1_ snapper probably contained a diverse mix of related individuals including full-siblings, half-siblings, and unrelated individuals. Approximately 80,000 larvae were reared in 5,000-L tanks held at 21°C and supplied with a filtered, UV-sterilised supply of incoming water. First grading of animals occurred on 1 May 2015, whereupon approximately 54,000 fish were retained for on-growing and supply. Grading was repeated every 2–6 weeks thereafter. Smaller (low-grade) animals were pooled separately from larger individuals but were retained at the same ration and water temperatures. Overstocking was prevented by grading fish into additional 5,000-L tanks as required. The water supply of the tanks was retained at temperatures above environmental (ambient) temperature, set to a minimum of 16°C. When fish were approx. 180 days old the cohort was graded and approximately 17,000 individuals were moved to a 50,000-L recirculating tank for further on-growing. An additional 10,000 high-graded (large) individuals were retained in two 5,000-L tanks, and approximately 12,000 small (low-grade) individuals were separated into additional 5,000-L tanks. All fish were retained on equivalent rations and at temperatures above ambient. Stocking densities were held below 25 kg/cm^3^ in all instances. Final grading of the cohort was performed in the week of 23 November 2015. Fish suitable for transport to the seapen (i.e. fish in good condition, uninjured, and of normal conformation) were graded to a minimum of 100-mm fork length and hand counted, yielding 21,891 individuals. Transport to PFR’s sea pen in Beatrix Bay (Pelorus, Marlborough) took place on 16 December 2015, when fish were either 319 or 312 days old.

On 26–27 April 2017 the snapper culture trial was ended following 17.5 months of on-growing. The mean size of snapper at harvest was 238-mm fork length, where sizes ranged between 139- and 301-mm fork lengths (Cook and Black 2017). Of these snapper, 200 of the smallest and 200 of the largest were selected by eye for this study, roughly falling within the 5% extremes of the size distribution (representing 5% of the smallest and 5% of the largest fish).

### DNA extraction and library preparation

Total genomic DNA was extracted from fin tissue samples collected from the 5% biggest (L) and the 5% smallest (S) individuals (200 for each size group), following the protocol described by Ashton *et al*. (2019). The concentration, purity and integrity of the extractions were assessed using Qubit (Life Technologies), NanoDrop (Thermo Scientific) and 2% agarose electrophoresis gels, respectively. The 188 samples with the best quality for each size group, plus eight replicates, were used for the preparation of 364 libraries using the reduced genome representation technique of ddRADseq (double-digest restriction site-associated DNA sequencing). We followed the protocol described by Peterson *et al*. (2012) with a few modifications as described by Sandoval-Castillo *et al*. (2018). For each sample, approximately 300 ng of DNA was digested with the restriction enzymes *Sbf*I and *Mse*I, and then ligated with forward and reverse adaptors, with forward adaptors including one of 96 individual barcodes designed in-house. Libraries were size selected for 250–800 bp fragments with a Pippin Prep (Sage Science), and then amplified using PCR. Individual libraries were mixed in equimolar concentration in four pools of 96 samples, and each pool was sequenced in an Illumina HiSeq 4000 lane (paired-end 150bp) at Novogene Co.

### Sequence quality checking and filtering

Sequence quality was checked using FASTQC 0.11 (Andrews 2011). Raw reads were then de-multiplexed into individual samples using ‘process_radtags’ from STACKS 1.5 (Catchen *et al*. 2013). Barcodes, rag-tags, bad quality base pairs (slide window 5, Q<25) and adapters were trimmed using TRIMMOMATIC 3.9 (Bolger *et al*. 2014). Reads with more than 5% “N’s”, average quality lower than 20, and length shorter than 70 bp were also removed. SNPs were called using the GATK pipeline, following recommendations by Van der Auwera *et al*. (2013). We used BOWTIE 2.3 (Langmead and Salzberg 2012) to align filtered reads to the chromosome-level Australian snapper genome (Ashton *et al*. 2019b;). We improved the alignments by locally realigning indels, then called SNPs using BCFtools 1.19 (Danecek *et al*. 2021). To reduce the number of false variants produced as inherent artefacts of the ddRAD sequencing approach (e.g. paralogous, sequencing errors, allele drop), several filtering steps were applied to the raw, called SNPs (Table S1) using VCFTOOLS 0.1.16 (Danecek *et al*. 2011). Firstly, we removed SNPs called in < 80% of the samples, with a minimum allele frequency < 3%, or with allele balance < 20 or > 80% on heterozygote genotypes. After removing indels, and to avoid sequencing or misalignment errors, we selected SNPs with ≥ 20% depth/quality ratio and ≥ 30 mapping quality (Li 2014). Additionally, to reduce paralogous loci and allelic dropout, we eliminated SNPs with too high or too low coverage (> mean depth plus twice the standard deviation, or ≤ 10X average depth per individual). Loci that did not conform to Hardy Weinberg Equilibrium in samples of both size groups, missing in more than 5% of samples of either of the size groups, or that showed discrepancy between two or more replicates, were also removed. For some analyses, we created an unlinked data set by pruning SNPs using a window size of 100 kb, a variance inflation factor of 5 and a R^2^ of 0.5 in PLINK 1.9 (Chang *et al*. 2015).

As recommended for GWAS analyses, we removed individual that deviate ±3 standard deviations from the average heterozygosity (Marees *et al*. 2018). Because inbreeding and relatedness can affect GWAS analyses, we calculated the individual inbreeding coefficient (F_IS_) in PLINK 1.9 and pairwise relatedness in VCFTOOLS using the unlinked SNP dataset. We then compared the average F_IS_ and relatedness of each size group using a non-parametric Mann Whitney U test. Age, sex, and population structure are factors that can potentially influence GWAS analyses, but given all fish used in this study were from the same population and age class and were immature at the time of the terminal sampling, these factors were considered not to bias the GWAS analyses.

With the aim of identifying SNPs involved in growth-related traits, we compared the allele frequencies between size groups using three different methods, each of which implement a different approach to control for relatedness and possible inbreeding. Firstly, based on identical-by-state pairwise genetic distances, we used complete linkage agglomerative clustering to create multidimensional scaling plots (MDS). Then, to control for family group substructure, we used MDS components as co-variables in a logistic regression implemented in Plink and calculated asymptotic *p* values to test for significance. This approach tests for odds of the minor allele frequency being associated with either phenotype (in this case large or small fish), when adjusting for potential confounders. Secondly, a pairwise kinship matrix was used as a co-variable in a linear polygenic mixed model. The significance of the model was calculated using the FAmily-based Score Test for Association (FASTA), all implemented in the R package GenABEL (Aulchenko *et al*. 2007). Thirdly, the same pairwise kinship matrix was used as a co-variable in a mixed linear model, but the significance was tested using a combination of a sequence kernel association test (SKAT) and a likelihood-ratio (LR) test, comparing a model with association against a model of non-association; these approaches were implemented using the R package rainbowR (Hamazaki and Iwata 2020). Like GenABEL’s model, rainbowR tests for a significant contribution of a polygenic component in the total genetic variance between size groups, when considering a random component and multiple covariance component effects. The main difference is implementation of LR in rainbowR, which is expected to better control for false positives (Hamazaki and Iwata 2020).

These analyses were based on single-SNP effects, but considering that gene function could be more commonly influenced by the small effects of several SNPs combined than the large effect of one SNP, haplotype-based approaches can improve the detection power of causal variance (Hamazaki and Iwata 2020; Sinclair-Waters *et al*. 2020). We therefore implemented two different methods to calculate haplotype blocks in rainbowR. One method considers pairs of variants within 200 kbp of each other that show high coefficients of linkage disequilibrium (D_lowerCI90_>0.7 and D_higherCI90_>0.98), as implemented in Plink (Chang *et al*. 2015). The other method uses a sliding window of 15 to cluster the closest 15 SNPs, implemented directly in rainbowR (Hamazaki and Iwata 2020). In addition to investigating the genetic architecture of growth in Australian snapper, we examined the proportion of size variance explained by each SNP and chromosome using the package bayesR (Moser *et al*. 2015) with default priors. This software estimates the genetic variance explained by each SNP and describes the genetic architecture of complex traits using Markov chain Monte Carlo and Bayesian mixture models. We calculated SNP-based heritability (h^2^_SNP_) as the proportion of phenotypic variance explained by all SNPs and considered this an approximation of the genome narrow-sense heritability (h^2^_g_). Accuracy of genomic prediction was evaluated using ten runs of bayesR, in each of which 80% of samples were randomly selected as training data and the remaining 20% was used as validation data. Since we used size as a categorical variable, the accuracy was measured in the validation sets as the beta coefficient of a logistic regression (predicted phenotype as a function of real phenotype) divided by the square root of heritability (h^2^_SNP_).

To explore the function of the candidate SNPs, we clustered SNPs in sliding windows of 10 kbp and extracted the sequence of each cluster from the reference genome. These sequences were then aligned to a Teleostei protein database obtained from UniProt (2021), using blastx 2.2.28 (Camacho *et al*. 2009) with a minimum alignment length of 30 amino acids, a similarity score of 50%, and an e-value threshold of 1×10-06. Enrichment of genetic functions on the putative candidate genes was explored using the gene ontology (GO) terms provided by UniProt and the R package topGO 2.26 (Alexa *et al*. 2016). Pathways and protein interaction analyses were performed using the web servers String 11 (Szklarczyk *et al*. 2021) and reactome 3.7 (Jassal *et al*. 2020).

## Results

Approximately 3.6 billion raw reads were obtained from the four Illumina lanes. After de-multiplexing, trimming and filtering, we retained over 3.5 million pairs of reads per sample, from which > 83% were successfully aligned to the snapper reference genome. From these, 1,273,017 variable sites were identified. After strict filtering, the dataset was reduced to 17,490 high-quality SNPs, which were evenly distributed among the 24 chromosomes of the reference genome (Table S2). After removing replicates and 12 samples (five large and eight small fish) owing to large amounts of missing data or extreme heterozygosity, 363 individuals (183 large and 180 small) were retained, with an average of 0.8% missing data.

The mean inbreeding coefficient was slightly higher in the small fish group (large=-0.1225 ± 0.1990; small= −0.0486 ± 0.1805, *p*<0.01). However, all individuals in both groups had inbreeding coefficients lower than 0.15, with the average per group lower than 0. As such, we considered the extent of inbreeding to be low in both groups. Average relatedness was not significantly different within compared with between the two size groups (large = 0.0089 ± 0.1320; small = 0.0098 ± 0.1170; large vs small = −0.0071 ± 0.1353; all *p* > 0.1).

The three single-SNP-based methods identified 31 candidate loci under selection between large and small fish. Of these nine, 11 and 12 were selected by Plink, GenABEL and rainbowR, respectively (Fig. 2). None of these was identified by two or more tests; however, these 31 candidates were in 10 chromosomes and one scaffold, and most of them (18 loci; 58%) were in physical proximity to one region of chromosome 16 (Fig. 2). The haplotype-based methods identified 77 SNPs in seven unique blocks as candidates, with three of these blocks identified by the two approaches (Fig. 3). The seven blocks were localized on five chromosomes and one scaffold, but more (three blocks; 43%) were found on chromosome 16 than on any other chromosome (Fig. 3). All these results suggest a key role of chromosome 16 on growth in the Australian snapper. This was concordant with the bayesR results, which found that the SNPs on chromosome 16 explained 16.4% of the size variance in our samples (Fig. 4, Table S2). Interestingly, this analysis suggested that a significant amount of growth variance in the Australian snapper can be explained by large effects of a relatively small number of SNPs (60 SNPs; Table S3). However, to explain as much as 76% of the phenotypic variation, small effects of many loci need to be considered. Based on the ten repetitions of cross-validation, the phenotype prediction accuracy was considerably high (74.4%), but expected given that the set of SNPs can explain most of the phenotypic variation in our data.

**Figure 2.**
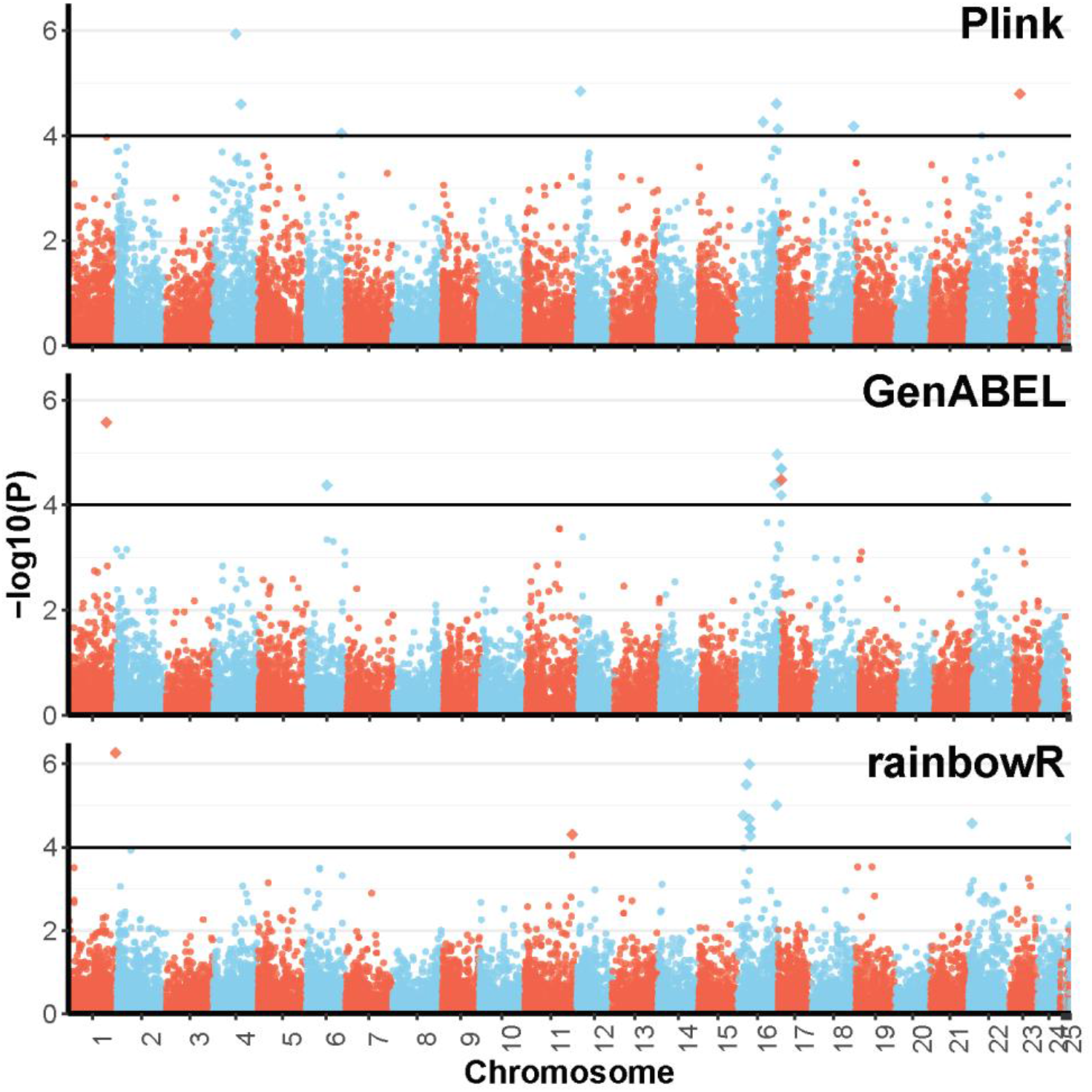
Manhattan plot of results from the three single-SNP-based genome wide association analyses of growth for 363 Australian snapper (*Chrysophrys auratus*) using 17,490 SNPs. SNPs are plotted according to their chromosomal position against their –log10 (*p*-value). Significant SNPs are the diamonds over the horizontal line (-log10(P)>4).

**Figure 3.**
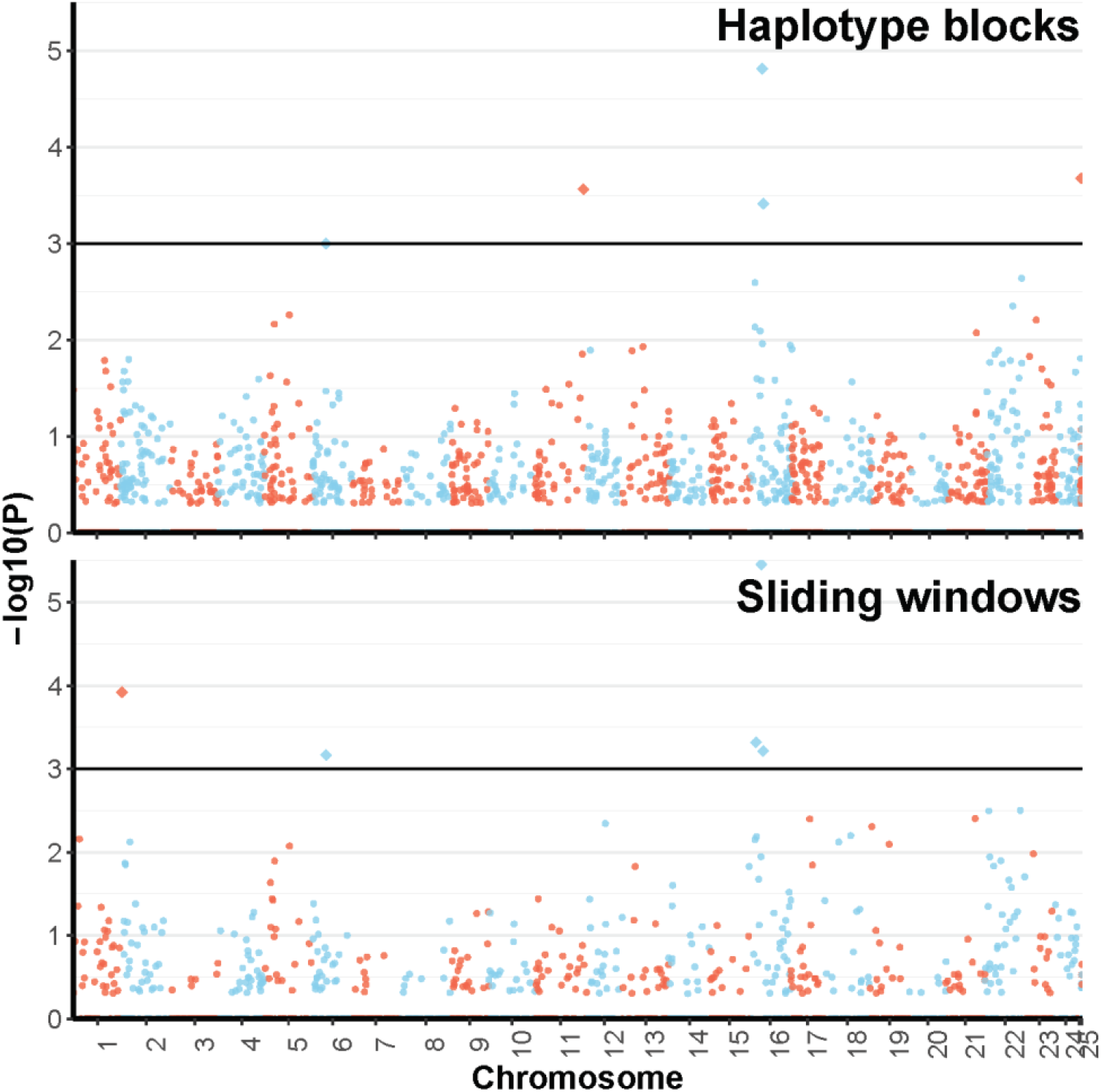
Manhattan plot of results from the two haplotype-based genome wide association analyses of growth on 363 Australian snapper (*Chrysophrys auratus*) using 17,490 SNPs in 1,337 blocks and 8,201 SNPs in 2,959 windows. Haplotypes are plotted according to their chromosomal position against their –log10 (*p*-value). Significant haplotypes are the diamonds over the horizontal line (-log10(P)>3).

**Figure 4.**
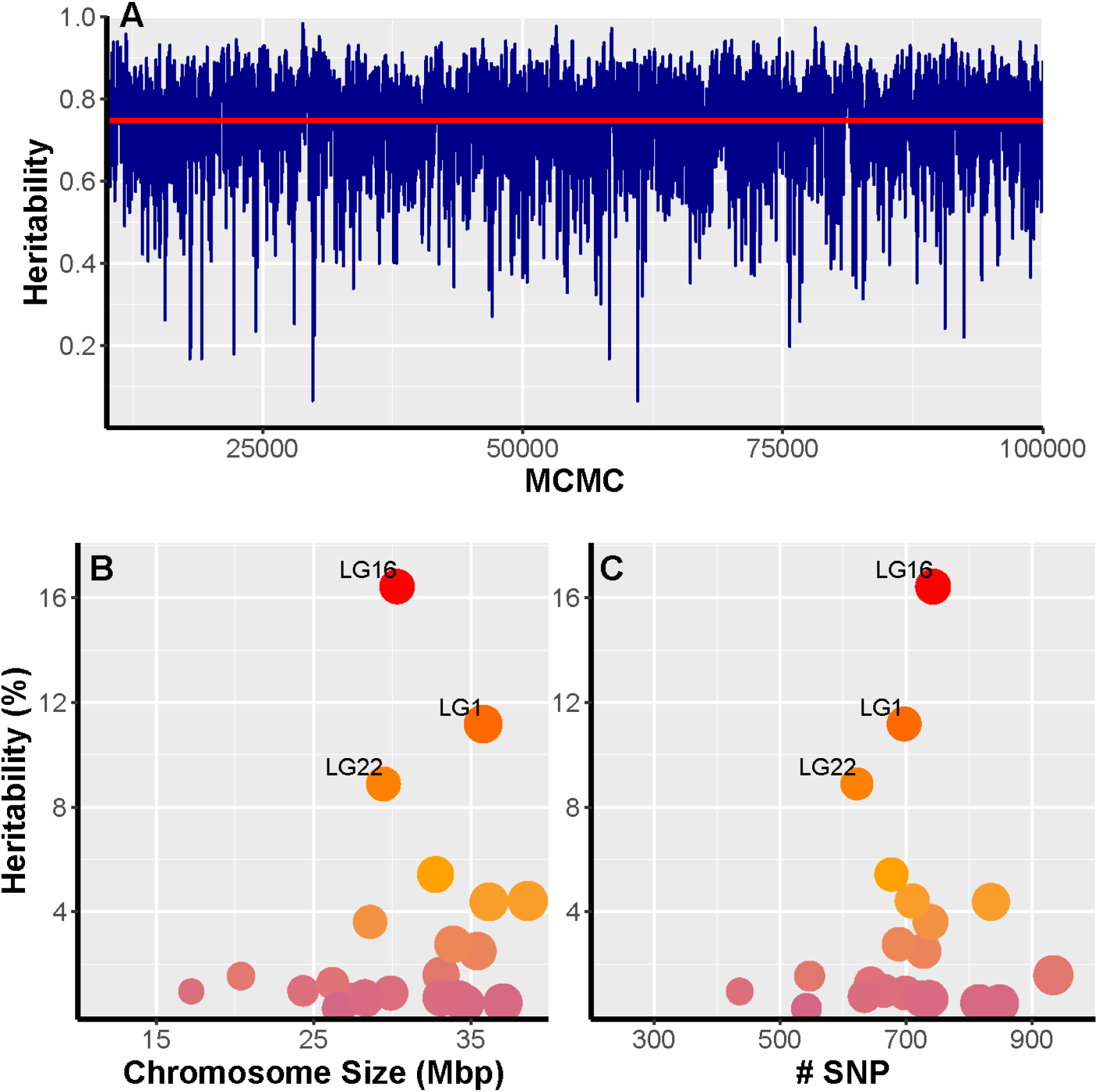
SNP-based heritability (h^2^_SNP_) of growth in Australian snapper (*Chrysophrys auratus*) estimated from Bayesian mixture models using 17,490 SNPs in 24 chromosomes. A) Convergence of MCMC sampling with average heritability (h^2^_SNP_=0.757) shown by the red horizontal line. Proportion of heritability or genetic variance of growth explained by each chromosome as a function of B) chromosome size in megabase pairs and C) number of SNPs per chromosome.

From both single-SNP and haplotype block approaches, 100 unique candidates were identified. We extracted 65 contigs of 10 kbp that included all unique candidate SNPs. Of these contigs, 57 were annotated to 51 proteins, where 16 SNPs were localized in exon regions. The proteins were assigned to 153 GO terms, but none of the functions or pathways was significantly enriched (p=0.18; results not shown). Despite this, most of these genes were involved in the metabolism of proteins, cell organization and tissue development, including five directly involved in growth (Table S4).

## Discussion

In this study we successfully identified 100 growth-related SNPs, of which 60 were located on chromosome 16. We also provide estimates of parameters related to the genomic architecture of growth, including heritability, and the number and effect sizes of SNPs contributing to size variation. These results demonstrate that the reduced genome representation SNPs, combined with appropriate bioinformatic tools, provide a cost-efficient approach to identify growth-related loci and to broadly describe complex trait genomic architectures. Our results provide crucial information to aquaculture breeding programmes about the targets of selection to enhance growth in this and related species. Knowledge about the genomic basis of growth also improves our general understanding of the impacts of anthropogenic stressors, such as climate change and extractive fisheries, on growth trait evolution in wild fisheries stocks.

### Genome architecture of a complex growth trait

Several studies have identified genes with large effects underlying simple Mendelian traits, but because of the statistical limitations of single locus GWAS approaches, they have been less successful with polygenetic quantitative traits like growth or size (Wellenreuther and Hansson 2016). Perhaps the most famous example of this is human height, where even with large samples sizes (~250 thousand people), less than 20% of variation can be explained using genome-wide markers (Yengo *et al*. 2018; Sohail *et al*. 2019). Akin to many other non-human studies, our study relied on a relatively small sample size, which reduced the power to detect loci of medium or small effect. However, by transforming the quantitative trait to a qualitative trait (big vs small) by comparing extreme phenotypes representing the 5% extremes of the phenotypic variation, and by using both haplotype-based and Bayesian mixture model approaches, we not only identified several growth-related loci, but also confirmed the polygenetic nature of the trait in the Australian snapper. As such, our study provides a clear example of how switching the focus away from single significant loci, to using information from all genotyped SNPs, can improve the statistical power to detect polygenic components of important phenotypic variants, including some regions with small but significant effects.

Our results suggest that a relatively small number of SNPs (<20K) is enough to explain most of the phenotypic variation related to growth. The overall heritability was generally higher than those reported in other studies (but see Valenza-Troubat *et al*. 2021). However, the 100 candidate SNPs explain similar phenotypic variation (4.4%) to those in other fish studies (0.1–19.0 %; Wu *et al*. 2019; Gong *et al*. 2021), including a quantitative trait loci analysis of the same species (Ashton *et al*. 2019b). As expected in complex traits like growth, individual candidate effects were small and multiple candidates were required to explain most of the phenotypic variation (Wellenreuther and Hansson 2016; Sinclair-Waters *et al*. 2020). However, only four candidates in the last 100 kbp of chromosome 16 could explain up to 3% of the total phenotypic variation. In addition, if we consider all SNPs in this chromosome, the explanatory power is elevated to over 16%. The variation explained by the markers on chromosome 16 is very high and could suggest physical linkage of genetic variation associated with growth. This is consistent with empirical examples of evolution of multiple linked variations that together modify the function of a gene or a complex of genes (Koshikawa *et al*. 2015; Kingman *et al*. 2021). This type of evolution promotes the formation of tightly linked haplotype blocks, allowing for the selection and inheritance of multiple sites with effect over several aspects of a complex trait, such as growth.

Haplotype-based evolution of polygenic regulation of growth could explain the lack of overlapping candidates from different analyses. The statistical approach of each method will capture loci with a particular degree of effect, and therefore only a fraction of the complex polygenic architecture. Such architecture would be more likely to include a mixture of loci with large, medium, small, pleiotropic, and synergic effects (Wellenreuther and Hansson 2016; Ashraf *et al*. 2020; Sinclair-Waters *et al*. 2020). Although the selected candidate SNPs differed between methods, all methods selected at least one candidate SNP within the last 5% of chromosome 16’s length. This chromosome region is broadly consistent with putative growth-related quantitative trait loci peaks identified for three growth traits in one-year-old Australian snapper reared in a land-based finfish facility (Fig. 5 in Ashton *et al*. 2019b). In that study, chromosome-level additive variance was not calculated; however, their results also suggested that chromosome 16 tended to explain most of the phenotypic variation. Here, neither chromosome length nor SNPs per chromosome was significantly correlated with the proportion of variation explained (Fig. 4). This suggests that the high relative contribution of chromosome 16 is not just an artefact of our reduced genome representation approach, but an important part of the genetic architecture of growth in Australian snapper. The results from this study being in part consistent with those by Ashton *et al*. (2019) also shows that the genomic basis of growth is comparable across different rearing environments (land-based finfish facilities vs a seapen). Chromosome 6 showed the second largest number of candidates, but the SNPs in this chromosome explained less than 4% of the phenotypic variation. In contrast, chromosomes 1 and 22 each explain over 8% of the phenotypic variation, despite chromosome 1 having a similar number of candidates and chromosome 22 having only a few candidates compared with chromosome 6. This is additional evidence for the very complex genomic architecture of growth, which is composed by a mixture of several loci with small effects detected by the haplotype approaches on chromosome 6, and a few loci with larger effects detected by single-SNP approaches, especially on chromosome 22. Thus, it is important to combine conceptually different approaches to extract information from all markers, to gain a better understanding of the genetic components of phenotypic variation (De Maturana *et al*. 2014; Akond *et al*. 2021; Gong *et al*. 2021).

### Candidate genes

The high-quality genome assembly made it possible for us to identify 51 genes surrounding the 100 SNP candidates (Table S4). Although most of these SNPs were in introns, several were in the proximity of, or inside, exons of the relevant candidate gene. Nevertheless, we are cautious about inferred associations. Previous studies have suggested several candidate genes and pathways for teleost growth-related traits (De-Santis and Jerry 2007; Johnston *et al*. 2011). Of these, the somatotropic axis is perhaps the most important, since it plays a central role in the regulation of metabolic and physiological processes involved in fish growth (De-Santis and Jerry 2007). Although the key genes in this axis are the growth hormone (Irving *et al*. 2021) and the insulin-like growth factors, several other factors, carriers, and receptors are implicated in this pathway. Several of our candidate SNPs were near the genes from the somatotropic axis, including the fibroblast growth factor 18 (FGF18) and the growth factor 5 (GDF5). Both genes are essential regulators of cell growth, cell differentiation, morphogenesis, tissue growth and tissue repair, especially during skeleton development (Jovelin *et al*. 2010; Nagayama *et al*. 2013), potentially affecting multiple growth traits. Concordantly, FGF18 has also been associated with differences in growth of the large yellow croaker, *Larimichthys crocea* (Zhou *et al*. 2019). Other two candidate genes detected, nucleophosmin 3 (NPM3) and transmembrane protein 132E (TMEM132E), are associated with the regulation of growth factors. NPM3 is predicted to be a histone chaperone protein that modulates replication of DNA and gene expression during tissue development (Wu *et al*. 2009). Because of its proximity, it co-activates with the fibroblast growth factor 8 gene, and both have fundamental roles during development, cell growth and proliferation in vertebrates (Kikuta *et al*. 2007). TMEM132E is a component of the cell membrane and is implicated in the regulation and transport of insulin-like growth factors. This gene is also involved in the development of the posterior lateral line (Li *et al*. 2015) which, as a mechano-sensory organ, has physiologically demanding growth requirements. This requires not just the addition of new cells but also the maintenance of correct functionality (Sapède *et al*. 2002).

Biosynthesis and retention of proteins is fundamental for tissue development, and the efficiency of these processes determines growth rate (Fraser and Rogers 2007). Seven of our candidates are directly involved in metabolism of proteins (Table S4), six of which are on chromosome 16. Of these, the prolyl 3-hydroxylase 1 (P3H1) and the serpin peptidase inhibitor clade H (SERPINH1) are notable in fishes for their implication in biosynthesis of collagen and assembly of collagen fibrils (Oecal *et al*. 2016; Tonelli *et al*. 2020). Collagen is essential to develop the structure and strength of bones, skin, muscle, and cartilage tissues, and therefore collagen biosynthesis is critical to maintain tissue integrity during somatic growth (Fraser and Rogers 2007). Moreover, collagen metabolic processes have been associated with changes in growth rates and responses to temperature increments in the Australian snapper (Wellenreuther *et al*. 2019). The variation in P3H1 and SERPINH1 could have a direct effect on the collagen metabolism efficiency and therefore on the growth rate of Australian snapper. Other important processes for tissue growth include cell proliferation, differentiation, migration, and adhesion. From our candidates, seven genes were associated with these cell processes (Table S4), including laminin subunit alpha 3 (LAMA3), related extracellular matrix protein 2 (FREM2) and periostin (OSF2). These three genes have a regulatory function in cell migration and adhesion, and they are involved in organ morphogenesis and tissue development of integuments, spinal cord, brain, eyes (Sztal *et al*. 2011), pharynx, fins (Carney *et al*. 2010) and the skeleton of fishes (Kessels *et al*. 2014). Finally, small difference in expression levels of specific genes have been shown to have significant effects on growth rate and metabolic efficiency (Fraser and Rogers 2007; Wellenreuther *et al*. 2019). Thus, variation in genes that act as transcript regularity factors can have important biological consequences. We detected several genes involved in gene regulation (Table S4); for example, the transcription cofactor vestigial-like protein 2 (VGLL2) regulates transcription by RNA polymerase II of genes during skeletal, muscle and neural morphogenesis (Johnson *et al*. 2011; Buono *et al*. 2021). Faster growth can be achieved by reducing energetic cost of protein metabolism, increasing transcription regulation adeptness, and cell migration and adhesion efficiency, which together can make a larger proportion of energy and cell proliferation available for growth. We consider all the above-described genes as of potential importance in the growth patterns of not only Australian snapper but perhaps of other teleosts.

### Applied relevance of genomic prediction of growth

The most important contribution of our results related to the genomic basis of growth is the potential to inform individual breeding values in terms of growth efficiency. This information has direct applications to selective breeding programmes of commercially important species, such as Australian snapper, but also for the breeding programmes of the gilthead sea bream (*Sparus aurata*) in the Mediterranean Sea, and the red sea bream (*Pagrus major*) in Japan. The increment of growth rate and its energetic efficiency can directly reduce production time and cost, leading to higher economic returns (Ye *et al*. 2017; Valenza-Troubat *et al*. 2021). However, the complexity and polygenetic nature of commonly involved traits has made it difficult to implement efficient genomic selective breeding programmes (Goddard and Hayes 2009; Ashton *et al*. 2019b). Here, we partially dissected the genetic architecture of growth and, in doing so, we have provided a set of SNPs that can assist genome selection of elite broodstock lines for commercial breeding programmes. Our results show that the use of reduced genome representation is sufficient to estimate breeding values (i.e. to predict phenotypes) with relative high accuracy even without pedigree information (Table S3; Fig. S1). Owing to the inclusion of most of the causal mutations, and decreased limitation due to linkage between SNPs and causal mutations, the use of whole genome resequencing (WGS) could increase predictive ability. However, empirical results comparing the use of WGS versus a set of SNPs show nil to marginal increase in genomic prediction accuracy (Lu *et al*. 2020; Yoshida and Yáñez 2021). This suggests WGS is unnecessary for this purpose and supports our suggestion that reduced genome representation provides accurate genomic prediction estimates to assist in the selection of elite broodstock lines. For systems where hundreds or thousands of samples may need to be evaluated over time, such as for Australian snapper aquaculture, this has significant cost implications. Recent advancements in the development of cost-effective genotyping technologies, such as SNP chips, are being developed for this, and related species (Montanari *et al*. in preparation), and will prove crucial in enabling the cost-effective assessment of commercially relevant SNPs.

Our data have the potential to determine individual breeding values, but this can be extended to the population level, with broader evolutionary and ecological implications. Estimating a genotype value and its frequency in a population could make it possible to predict how the population will respond to a future environmental event in respect to the phenotypic trait in question (Hunter *et al*. 2021; McGaugh *et al*. 2021). This can be an important tool in the re-stocking of heavily affected populations, either by climatic change or by overfishing. There are several factors affecting growth in marine organisms, but one of the most important abiotic factors is temperature (Besson *et al*. 2016; Boltaña *et al*. 2017; Wellenreuther *et al*. 2019). Furthermore, climate warming and associated environmental changes are considered great threats to the future of both wild fish biodiversity and global fisheries (Free *et al*. 2020). The most direct, immediate, and common fish responses to climate change are reflected in modifications to the growth rate (Rountrey *et al*. 2014; Ong *et al*. 2015; Huang *et al*. 2021). Although warmer sea surface temperatures are likely to increase the growth rates of most fishes, for individuals and species living closer to thermal optima, warmer waters might decrease growth rates (Neuheimer *et al*. 2011; Huang *et al*. 2021). In addition, many fish populations are under heavy exploitation, and there is strong evidence of selective harvesting gradually reducing the genetic potential for somatic growth in the population (Enberg *et al*. 2009; Denechaud *et al*. 2020), and thus magnifying climatic change impacts (Morrongiello *et al*. 2021; Wootton *et al*. 2021). Small changes in growth rates within a population can not only influence individual fitness but can also cause long-lasting shifts in population characteristics and demographic dynamics, including a reduction in fecundity, survival, and recruitment rates (Lorenzen 2016; Denechaud *et al*. 2020). For most marine fishes, mortality is considerably higher during early life stages. At that time, individual phenotypes influence the probability of survival, and both field and laboratory research have shown this effect to be size dependent (Johnson *et al*. 2014). Moreover, reproductive success is often broken into two components: reproductive potential and offspring survival, and in many marine fish species both components are strongly related to body size. Because female fish retain their oocytes internally during their development, maximum reproductive output will be subject to body size constraints (Lambert 2008; Ohlberger *et al*. 2020). Alternatively, reduced body size can result in small eggs that maximise the number produced, but with a significant reduction of offspring survival (Einum and Fleming 2000; Wootton *et al*. 2021). Since large organisms have relatively high survival probabilities and reproductive success, it is expected that by predicting the size composition of a population, we can determine its average survival and recruitment dynamics (Garrido *et al*. 2015). In other words, introducing size-enhanced genotypes to wild populations to modify natural size distributions could in turn increase the resilience to both climatic change and harvesting. Thus, size genomic prediction can be a valuable tool for fishery and conservation management, not only for Australian snapper, but also for other marine fishes.

Using breeding values to predict population response has potential complications, such as failure to incorporate the complexity of factors and uncertainty involved in trait measurement (Hadfield *et al*. 2010). A wide range of factors affects growth in marine fishes, such as temperature (Boltaña *et al*. 2017; Wellenreuther *et al*. 2019), nutritional state (Escalante-Rojas et al 2020) and intra- as well as inter-species interactions (Mallard *et al*. 2020; Korman *et al*. 2021). Although these factors can induce phenotypic plastic changes to growth, there are also important genetic components in response to these factors (Wellenreuther *et al*. 2019; Escalante-Rojas *et al*. 2020). Adaptive variation in our candidate genes that regulate expression and synthesis of stress-related proteins, such as SERPINH1 and PRKAG2 (Wang *et al*. 2016; Causey *et al*. 2019), can potentially constrain these responses in Australian snapper and, therefore, the effects of external factors. In addition, Bayesian approaches, such as the one used here, can integrate the effects of unknown factors and uncertainty involved in genomic predictions in an efficient way (Ashraf *et al*. 2020). This suggests that our genomic predictions provide a reliable way to measure individual breeding values of Australian snapper from both wild and captive populations.

This study confirms that, despite the complex polygenetic architecture of growth in Australian snapper, reduced genome representation combined with a mix of bioinformatic approaches can detect candidate genes relevant to a quantitative trait. This opens the possibility of using Bayesian genomic prediction frameworks to measure individual breeding values for growth rates. This information is expected to assist the selective breeding programmes for this and related species, and can be used to provide insights into growth changes experienced by wild populations following the exposure of anthrophonic stressors, such as climate change and fishing.

## Acknowledgements

We would like to acknowledge the PFR staff who assisted with the breeding and husbandry operation of the snapper populations, in particular Warren Fantham, who oversees the rearing of finfish larvae, and Therese Wells, who manages the post-juvenile trevally husbandry. This research was funded through the MBIE Endeavour Programme “Accelerated breeding for enhanced seafood production” (#C11×1603) to M.W. and by the Australian Research Council (LP180100756) to L.B.B.

## Data availability statement

As the genomic data of this species are from a taonga and thus culturally important species in Aotearoa New Zealand, the data have been deposited in a managed repository that controls access. Raw and analyzed data are available through the Genomics Aotearoa data repository at https://repo.data.nesi.org.nz/. This was done to recognise Māori as important partners in science and innovation and as inter-generational guardians of significant natural resources and indigenous knowledge.

---

**Table S1.**
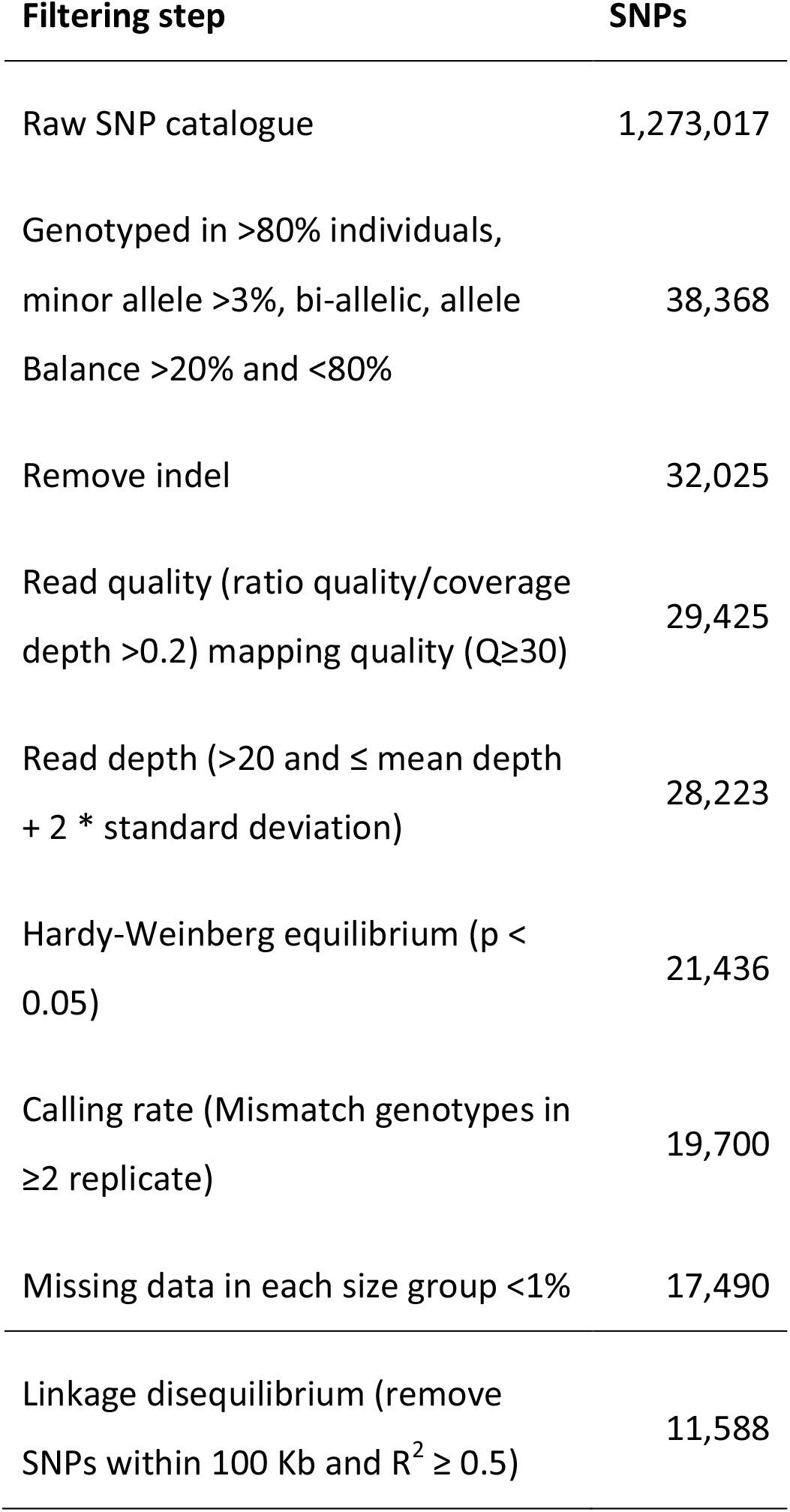
Filtering steps and number of SNPs retained for the two groups (small and large) of Australian snapper (*Chrysophrys auratus*).

**Table S2.** Chromosome size, SNPs per chromosome (bp = base pairs), and SNP-based heritability (h^2^_SNP_) per chromosome for Australian snapper (*Chrysophrys auratus*)

| Chr | bp | #SNP | SNP/100Kbp | $h^2_{\text{SNP}}$ |
| --- | --- | --- | --- | --- |
| LG1 | 35,796,517 | 697 | 1.95 | 11.18 |
| LG2 | 36,161,541 | 835 | 2.31 | 4.38 |
| LG3 | 34,661,116 | 849 | 2.45 | 0.49 |
| LG4 | 33,120,650 | 934 | 2.82 | 1.58 |
| LG5 | 35,422,278 | 728 | 2.06 | 2.50 |
| LG6 | 28,599,003 | 739 | 2.58 | 3.61 |
| LG7 | 34,240,341 | 725 | 2.12 | 0.66 |
| LG8 | 37,046,140 | 816 | 2.20 | 0.52 |
| LG9 | 27,277,866 | 739 | 2.71 | 0.66 |
| LG10 | 33,112,362 | 737 | 2.23 | 0.71 |
| LG11 | 38,628,475 | 710 | 1.84 | 4.41 |
| LG12 | 26,212,225 | 644 | 2.46 | 1.27 |
| LG13 | 33,878,147 | 689 | 2.03 | 2.76 |
| LG14 | 29,913,436 | 699 | 2.34 | 0.90 |
| LG15 | 28,243,467 | 723 | 2.56 | 0.70 |
| LG16 | 30,315,390 | 743 | 2.45 | 16.43 |
| LG17 | 24,333,991 | 665 | 2.73 | 0.98 |
| LG18 | 32,765,680 | 677 | 2.07 | 5.43 |
| LG19 | 28,261,070 | 634 | 2.24 | 0.77 |
| LG20 | 26,584,312 | 542 | 2.04 | 0.30 |
| LG21 | 28,350,763 | 729 | 2.57 | 0.64 |
| LG22 | 29,453,248 | 622 | 2.11 | 8.89 |
| LG23 | 20,385,525 | 547 | 2.68 | 1.54 |
| LG24 | 17,221,635 | 436 | 2.53 | 0.95 |
| LG25 | 3,820,059 | 40 | 1.05 | 0.03 |

**Table S3.**
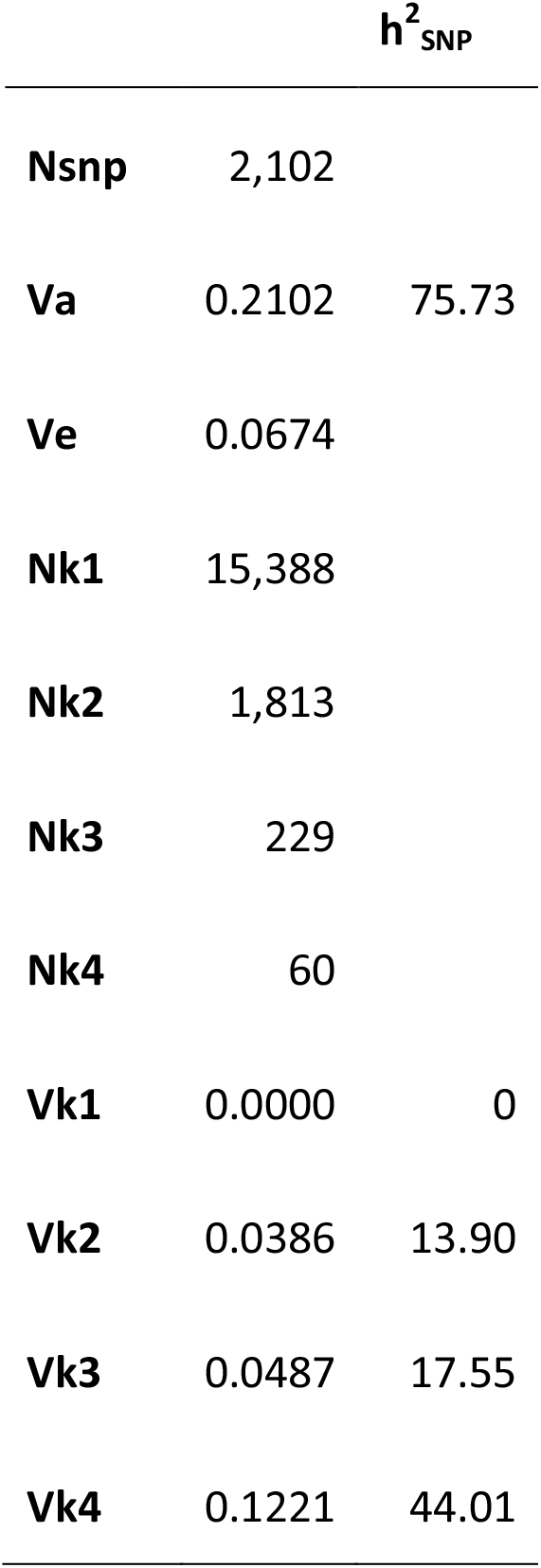
Model summary of bayesR results on the genomic architecture of growth of Australian snapper (*Chrysophrys auratus*). NSNP = number of SNPs use in the model to explain the variation, Vg = Genetic variance explained by SNPs in the model, Ve = residual variance, Nk[1-4] = number of SNPs in mixture component 1-4, Vk[1-4] = SNP effects in mixture component 1-4. Bayesian model assumes a variance in each mixture component of 1 = 0, 2 =0.0001, 3=0.001, 4=0.01. SNP-based heritability (h^2^_SNP_) explained by the whole model and each of its mixture components.

**Figure S1.**
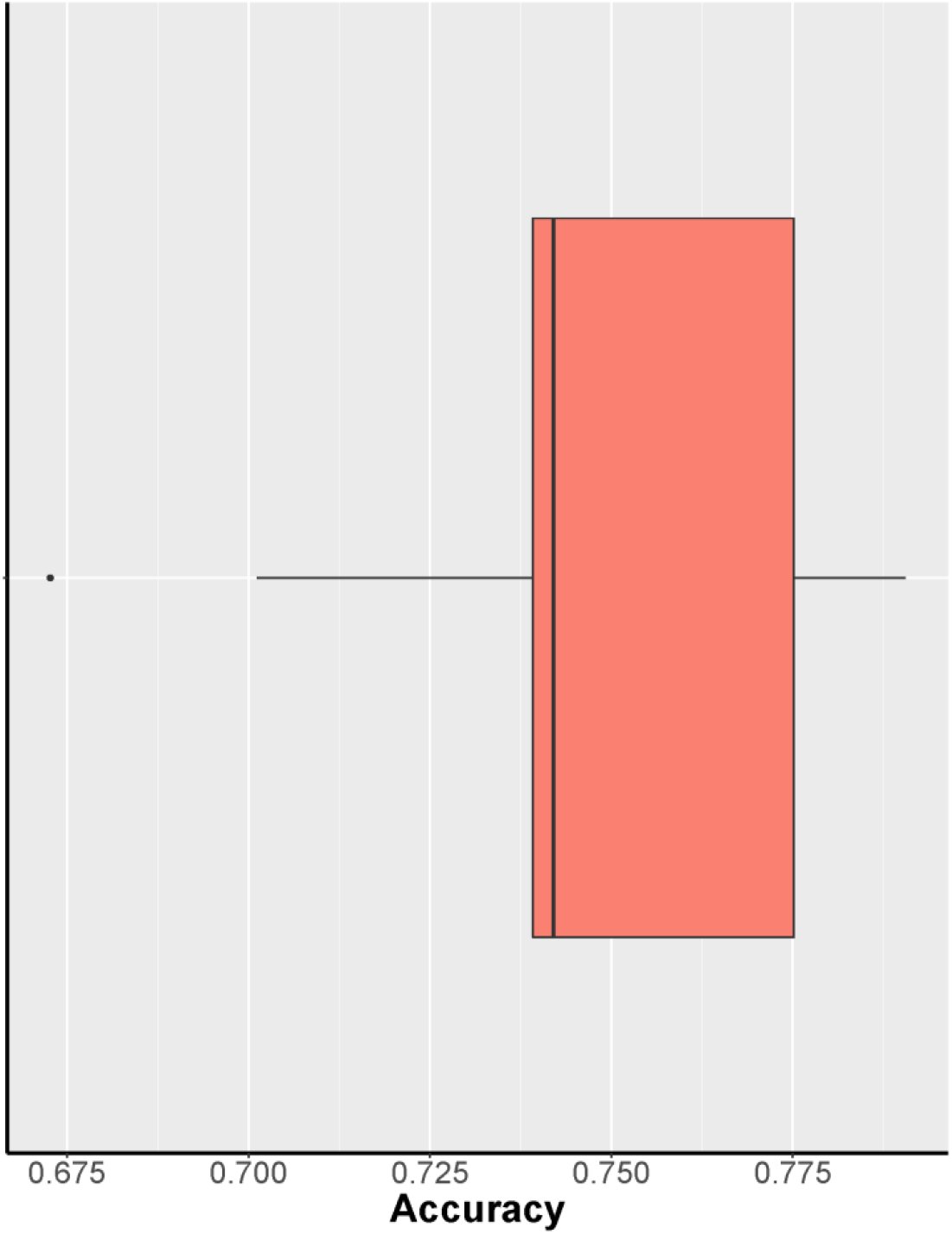
Boxplot showing the mean (0.744) and distribution of the accuracy of phenotype (small or large) prediction base on bayesR results using 17,490 SNPs for 363 Australian snapper (*Chrysophrys auratus*). The plot is based on 10 replicates with randomly selected 80% of the samples as training data and 20% of the samples as validation data.

